# Conjunctive Representations that Integrate Stimuli, Responses, and Rules are Critical for Action Selection

**DOI:** 10.1101/835652

**Authors:** Atsushi Kikumoto, Ulrich Mayr

**Author notes:** Corresponding Author: Ulrich Mayr, Department of Psychology, University of Oregon, Eugene, OR 97403, Fax: -4911.

## Abstract

People can use abstract rules to flexibly configure and select actions for specific situations. Yet how exactly rules shape actions towards specific sensory and/or motor requirements remains unclear. One possibility is that rules become integrated with sensory/response features in a non-linear, conjunctive manner (e.g., event files; Hommel, 1998) to drive rule-guided action selection. To dynamically track such conjunctive representations during action selection, we applied a time-resolved representational similarity analysis to the spectral-temporal profiles of the EEG signal, while participants selected actions based on varying rules. Across two experiments, we found that action selection engages conjunctive representations binding action rules to specific sensory/motor settings throughout the entire selection period. The strength of conjunctions was the most important predictor of trial-by-trial variability in response times (RTs) and was closely, and selectively, related to an important behavioral indicator of event files—the partial-overlap priming pattern. Thus, conjunctive representations were functionally dissociated from their constituent action features and play a critical role during flexible selection of action.

---

Flexible, goal-directed action requires the use of abstract rules that can be applied to a range of specific situations. However, we know little about how abstract rule representations connect with lower-level sensory or response representations, as a specific action is planned and executed. In traditional stage-based processing models, information flows from sensory to response in a cascade of relatively independent representations ^1–5^, that are specified by the relevant action rule.^6^ An alternative view is the idea of a common representational space in which all action-relevant features (e.g., sensory, motor, and even abstract action rules) are combined into highly integrated, conjunctive representations, sometimes referred to as *event files* or *task files*.^7–10^ By tying all relevant features together into a common, integrated representation, a specific action becomes executable. Therefore, these representations are a critical condition for successful action control and selection.

Once formed, however, an event file can also get in the way of subsequent actions, as indicated by a characteristic pattern of priming effects.^11^ Specifically, when consecutive trials require event files that share either all or none of the constituent features, actions are executed relatively fast. However, when only some, but not all features overlap across trials, then response-times or errors increase, a pattern that event-file theory explains as the cost of “unbinding” the overlapping features from the no-longer needed event file. Such partial-overlap costs emerge even when complete S-R associations repeat across trials while the abstract rule changes, indicating that just like any sensory or response features, rules can become part of event files.^10^

The partial-overlap pattern is currently the key empirical indicator of event files. However, because it is an *aftereffect* of event-file formation, this pattern provides no information about how conjunctive representations behave during response selection and whether or not they are indeed a critical precursor of successful action. Moreover, partial-overlap costs can also be explained by alternative models that do not assume integration between different codes during action selection. For example, Kleinsorge & Heuer^12, 13^ proposed a strict hierarchical separation between the level of rules and the level of stimulus/response selection. The pattern of partial-overlap costs arises from the assumption that a switch of action codes on the highest level (i.e., rules) propagates down to the lower levels (i.e., stimulus-response codes). As a result, when only the rule changes, but the response stays constant, the now inappropriate lower-level specification will have to be reverted, leading to performance costs.

To test the event file model against accounts that do not assume integration of action-relevant features as a critical step during action selection, it is important to directly track the multiple representations that could concurrently become active during action selection––including potential conjunctions between stimuli, responses, and even action rules. In the current study, we used the EEG signal to decode information about action-relevant representations in a time-resolved manner ^14–16^ via representational similarity analysis (RSA) ^17, 18^ as participants selected responses to location stimuli on the basis of randomly cued, spatial transformation rules^10^ (Fig.1ab). Experiment 1 allowed us to decode conjunctions that were specific for particular rules, but without differentiating between S-R conjunctions and conjunctions that also integrated abstract rules (i.e., rule-S-R conjunctions). In Experiment 2, we replicated all major results from Experiment 1, but also used an expanded task space that allowed us to test whether or not abstract rules can become integrated into conjunction representations. Across both experiments, we found strong evidence for conjunctive representations--including rule-S-R conjunction in Experiment 2. Consistent with predictions from event-file theory, conjunctions were robust and unique predictors of variability in performance, and were related to the pattern of partial-overlap priming costs.

## Results

### Experiment 1

#### Behavior

For all analyses, error-trials, post-error trials, and trials in which RTs were larger than 99.5 percentile of the RT distribution were excluded. Consistent with previous work^10^, we observed partial-overlap costs in RTs and errors as a function of the different trial-to-trial transitions (Fig. 2): When the rule, the stimulus, and thus also the response repeated or when all changed, responses were fast and accurate, whereas costs emerged in the case of partial updates of either rules or stimuli/responses (for statistical analysis, see Supplementary Table 1).

**Fig. 1.**
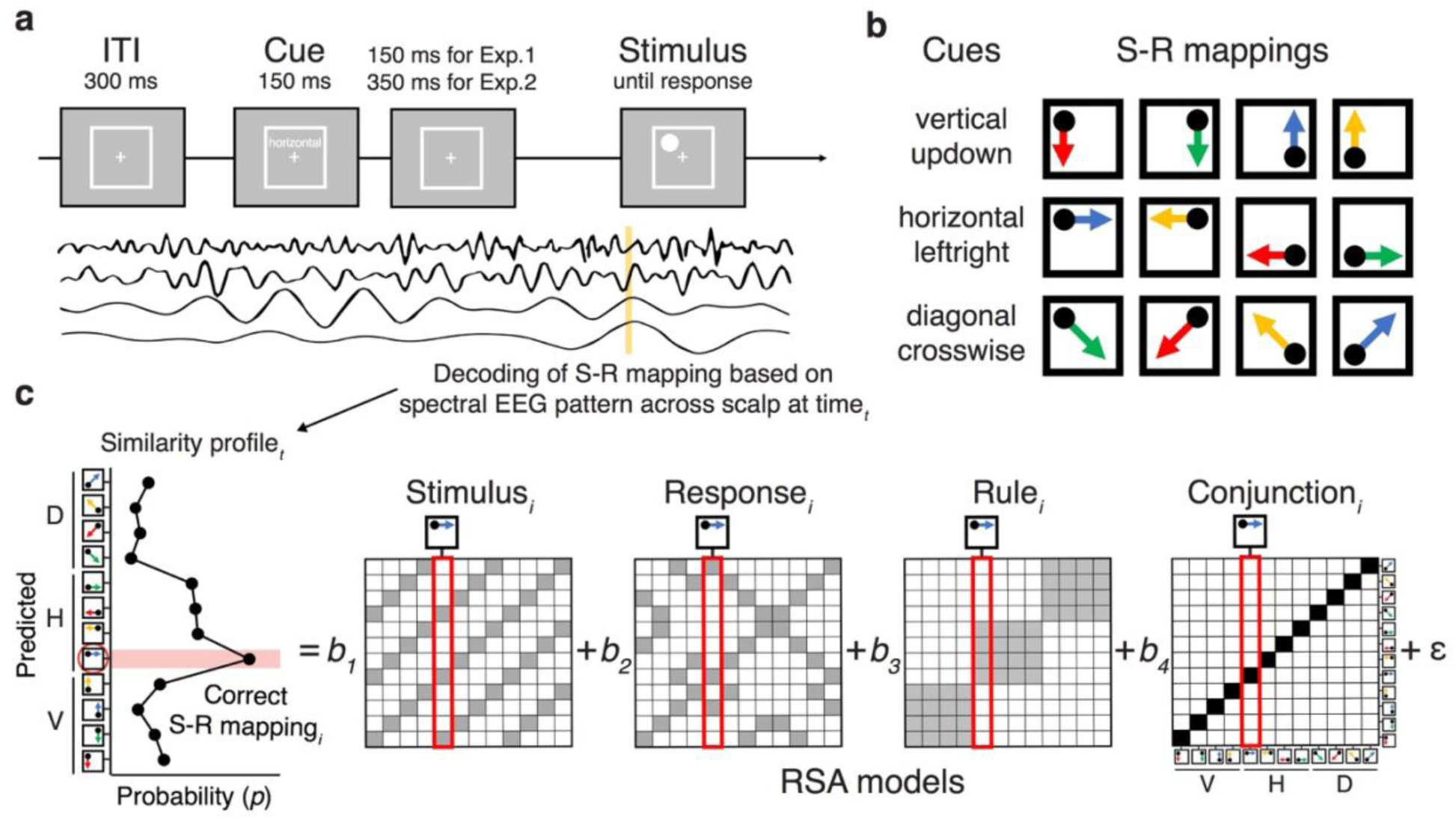
**a**, Sequence of trial events in the rule-selection task for both Experiment 1 and 2. **b**, Spatial translation rules mapping specific stimuli to responses in Experiment 1. Two different cue words were used for each rule. **c**: Schematic steps of the representational similarity analysis. The raw EEG signal was decomposed into frequency-band specific activity via time-frequency analysis (see *EEG recordings and preprocessing* and *Time-Frequency Analysis*). For each sample time (*t*), a scalp-distributed pattern of EEG power was used to decode the specific rule/stimulus/response configuration of a given trial, producing a set of classification probabilities for each of the possible configurations. The profile of classification probabilities reflects the similarity structure of the underlying representations, where similar action constellations are more likely to be confused. The idealized profile of classification probabilities shows an example where a unique conjunction and rule information is expressed (peak at the correct S-R mapping*i* and confusion to other instances with the same rule). For each trial and timepoint, the profile of classification probabilities is simultaneously regressed onto model vectors as predictors that reflect the different, possible representations. In each matrix of model vectors, the x-axis corresponds to the correct constellation for the decoder to pick, and the y-axis shows all possible constellation. The shading of squares indicates the predicted classification probabilities (darker shading means higher probabilities). The coefficients associated with each predictor (i.e., *t-*values) reflect the unique variance explained by each of the constituent features and their conjunction.

**Fig. 2.**
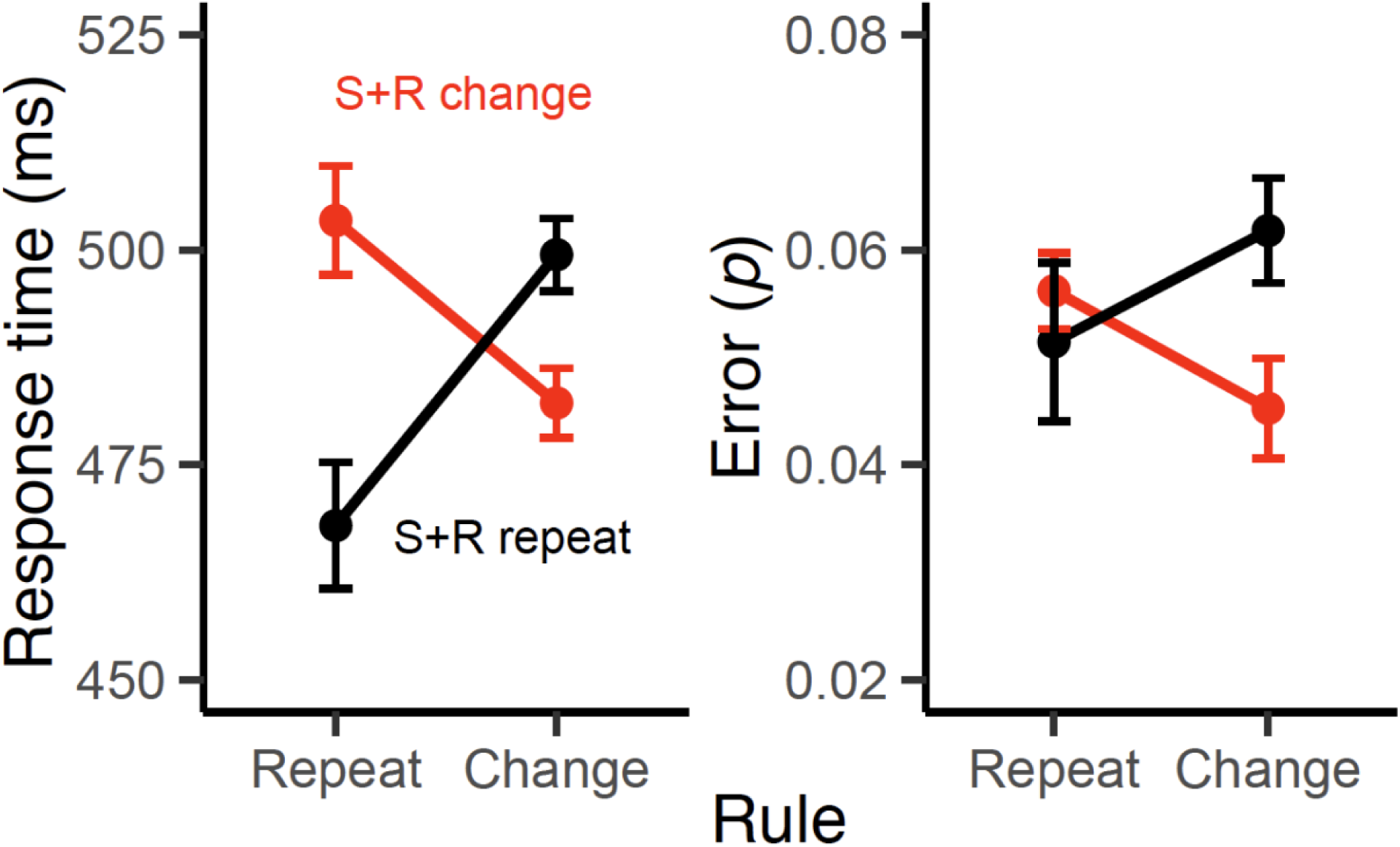
Mean response times (RTs) and errors for Experiments 1 as a function of rule repetition/change factor and the stimulus-response repetition/change factor. Error bars specify 95% within-subject confidence intervals.

#### Tracking Representational Dynamics

While the pattern of RTs and errors is consistent with predictions from the event-file model, by itself it is not sufficient to draw strong inferences about the role of conjunctive representations during action selection. Fig. 3a shows the results of the time-resolved RSA performed on the level of single trials. Consistent with previous results, the cascade of decoded representations unfolds in a manner that is consistent with the expected flow of information: The rule is activated during the pre-stimulus phase, followed by a strong expression of the stimulus, and finally by the response.^16, 19^ Critically, the conjunctive representation can be decoded during the entire post-stimulus period (Fig. 3a), and clearly peaks before response representations fully develops (Supplementary Fig. 7). These effects were significant even though we accounted for subject-specific differences in RTs between action constellations and therefore cannot be explained in terms of unspecific difficulty differences between action constellations (Supplementary Fig. 1).

**Fig 3.**
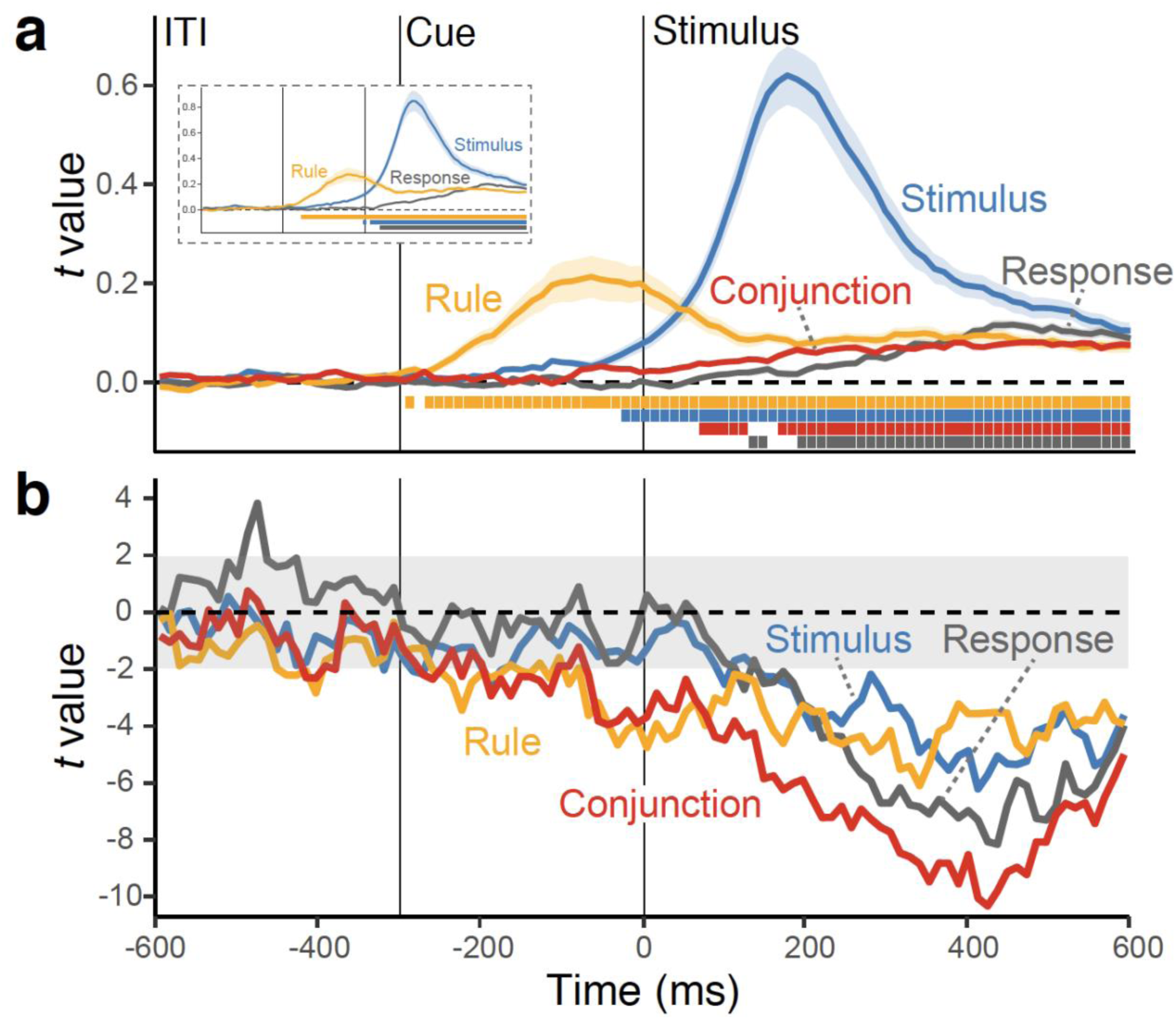
**a**, Average, single-trial *t*-values associated with each of the basic features and their conjunction derived from the RSA analysis (see Fig. 1c). Shaded regions specify the standard error around the mean. The colored squares at the bottom of the figure denote the significant time points using a non-parametric permutation test. The insert shows RSA fit scores when the conjunction was not included as predictor in the analysis. **b**, Time-course of *t* values from multilevel, linear models predicting the variability in trial-to-trial RTs (the “impact” of representations on behavior), using RSA scores of all features as simultaneous predictors.

To test the prediction from event-file theory that conjunction representations are critical for action selection, we regressed trial-to-trial variation in RTs onto the strength of each expressed representation. Using multilevel modeling, we performed these analyses for each time-point and with all predictors entered simultaneously. The resulting “impact-trajectories” are shown in Fig. 3b; statistical results for a-priori selected time intervals are summarized in Table 1. Note that negative *t*-values indicating that stronger representations lead to faster responding. Consistent with the prediction from event-file theory, the conjunctive representation was the most dominant predictor of performance. Combined, these results indicate that conjunctive representations emerge during response-selection, concurrently with the representations of constituent features, and predict upcoming behavior over and above the influence of the constituent representations.

**Table 1.** Predicting trial-by-trial RTs using the strength of decoded representations.

| Decoded variable | Pre-stimulus |  | Early Post-stimulus |  | Late Post-stimulus |  |
| --- | --- | --- | --- | --- | --- | --- |
|  | *b* (*se*) | *t*-value | *b* (*se*) | *t*-value | *b* (*se*) | *t*-value |
| Rule | -.021 (.005) | -3.89 | -.004 (.007) | -5.71 | -.037 (.007) | -4.97 |
| Conjunction |  |  | -.061 (.008) | -7.23 | -.093 (.011) | -8.07 |
| Stimulus |  |  | -.014 (.008) | -1.82 | -.041 (.012) | -3.49 |
| Response |  |  | -.027 (.006) | -4.12 | -.068 (.012) | -5.81 |

#### Conjunctive Representations and Partial-Overlap Costs

In order to directly connect the EEG-decoded conjunctive representations with event-files, we examined whether and how these representations relate to the partial-overlap priming pattern. As Fig. 4a shows, the strength of decoded conjunctions expresses the partial-overlap pattern. Conjunctive representations were particularly strong exactly in those transitions in which RTs were fast (i.e., when either everything repeats or everything changes, see Fig. 2). Conjunctive representations showed the partial-overlap pattern in the correct direction during early post-stimulus phase, *b*=-.024, *SE*=.010, *t*(20)=-2.58, but not in the late post-stimulus phase, *b*=-.004, *SE*=.010, *t*(20)=-.39, and none of the constituent features showed the critical interaction pattern, all *t*s(20)>-.21.

**Fig. 4.**
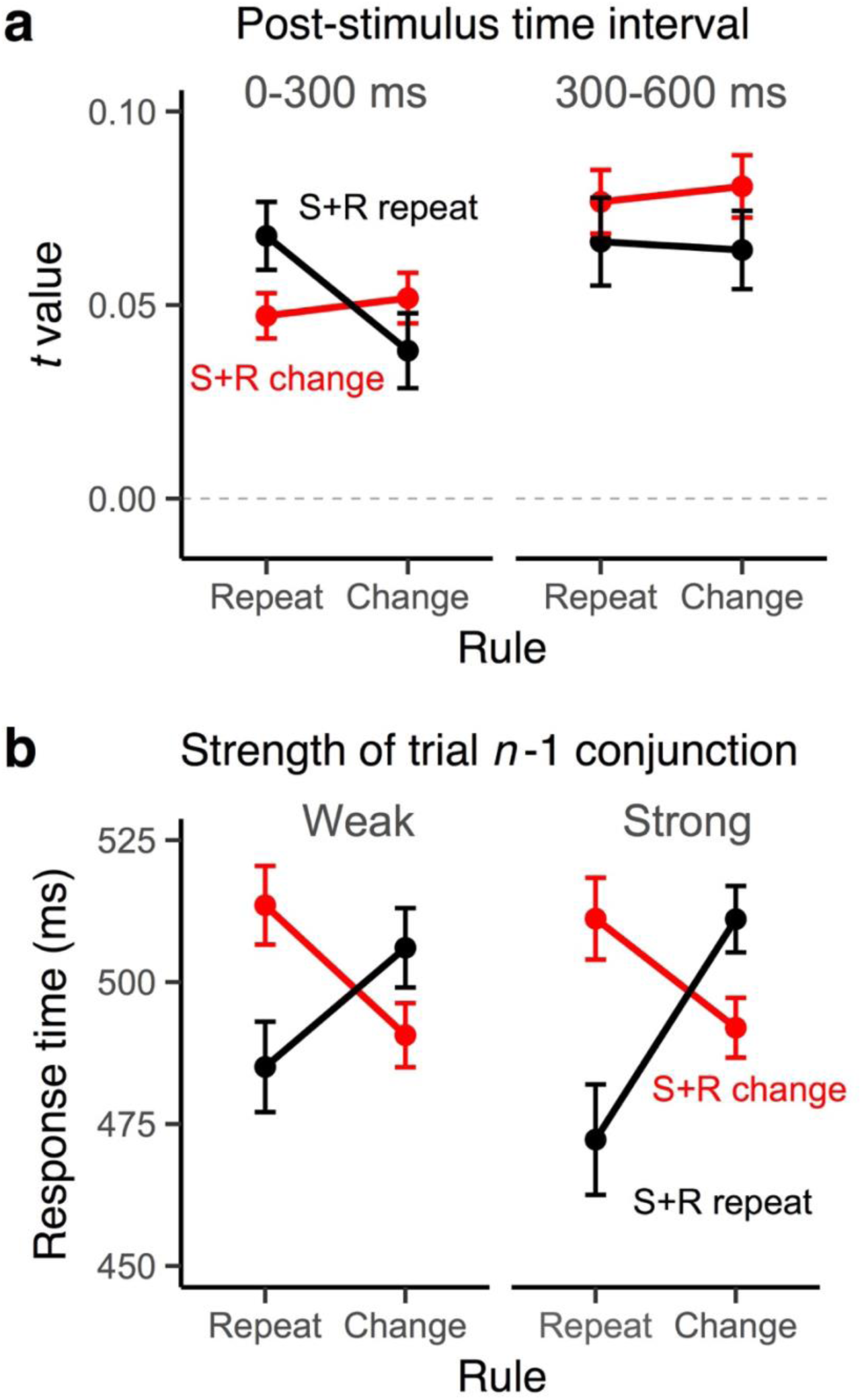
**a**, Average RSA scores of the conjunction model as a function of rule repetition/change and the stimulus-response repetition/change factors for two the early (0-300 ms) and the late (300-600 ms) periods in the post-stimulus interval **b**, Modulation of partial-overlap costs on RTs in trial *n* as a function of the strength of conjunction codes (median split) in trial *n* −1. Error bars specify 95% within-subject confidence intervals.

Another important prediction that can be derived from the event-file model is that strong conjunctions should be particularly difficult to “unbind” on the following trial. Thus, the strength of conjunctions on trial *n*-1 should predict partial-overlap costs on trial *n*. Our results, shown in Fig. 4b, confirm this prediction: A stronger conjunctive representation in *n*-1 trial, late in the selection period led to a greater RT partial-overlap costs on the next trial, *b*=-.025, *SE*=.011, *t*(20)=-2.25. Again, this pattern was unique for conjunction representations. None of the constituent features had comparable effects on next-trial performance, all *t*(20)>-.05. Taken together, the behavior of decoded conjunctive representations is highly consistent with predictions from the event-file model.

### Experiment 2

The results in Experiment 1 suggest that action selection recruits conjunctive representations and that the strength of these representations is predictive of trial-to-trial performance, as postulated by the event file perspective^8, 11, 20^. Yet, because each action rule specified a unique set of S-R links, the observed conjunctive representations could consist of any combinations of the rule and/or stimulus/response features—leaving it ambiguous to what degree abstract rules were integrated into conjunctive representations. Thus, in Experiment 2, we attempted to tease apart two different types of conjunctions: (1) rule-independent conjunctions between stimuli and responses (S-R conjunctions) and (2) rule-specific (rule-S-R) conjunctions that integrate abstract action rules with S-R links. To this end, we introduced four action rules (i.e., vertical, horizontal, clockwise, and a counterclockwise; Fig. 5a), which allowed S-R conjunctions that shared the same S-R links, but different abstract rules (e.g., a dot at the top-left corner requires a bottom-left response using either the vertical rule or the counterclockwise rule).^10^ The inclusion of such same-S-R pairs allows dissociating conjunctions that did integrate rules (rule-S-R) from rule-unspecific conjunctions (S-R).

**Fig. 5.**
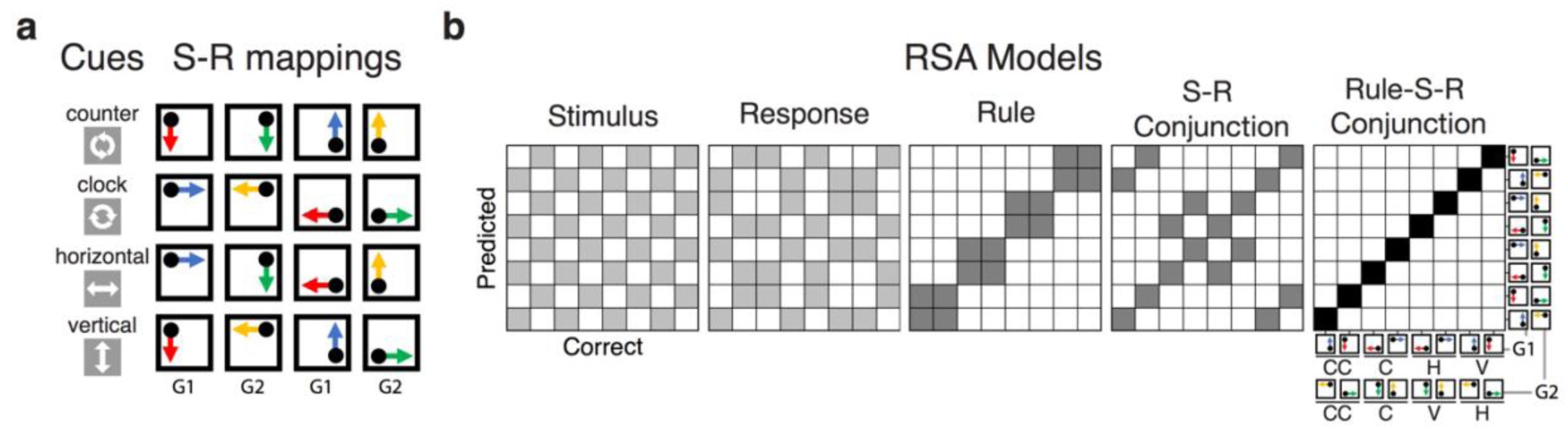
**a**, Spatial translation rules mapping specific stimuli to responses in Experiment 2. Either words or symbols were used as cues for each rule. S-R mappings were divided into two groups: G1 group (cases where a dot appeared at the top-left or bottom-right corner) and G2 group (cases where a dot appeared at the top-right or bottom-left corner) for the decoding analysis. **b**, Models for representational similarity analysis (RSA) in Experiment 2. The S-R conjunction model assumes a similar pattern for the specific combination of the stimulus and response irrespective of rules. The rule-S-R conjunction model expects a unique pattern for the configuration of each rule/stimulus/response combination. To completely orthogonalize action features, RSA was performed separately for G1 and G2 subsets of S-R mappings, requiring identical 8 x 8 model matrixes for each group. Analyses were performed separately within each subset and coefficients were averaged within subjects.

#### Behavior

The same trial-exclusion criteria as in Experiment 1 were used for all analyses in Experiment 2. We replicate the partial-overlap costs on RTs and errors from Experiment 1 and a previous report using the same paradigm ^10^ (Fig. 6): Critically, repetition of rule-S-SR settings produced RT and error benefits, whereas any partial updates (including S-R repetitions) generated costs. We focused on trials in which both stimulus and response-features covary (i.e., complete S-R changes or repeats) as a function of switching of rules because they provide a direct test for potential differences in the rule-specific and rule-independent conjunctions (for statistical analysis, see Supplementary Table 2).

**Fig. 6.**
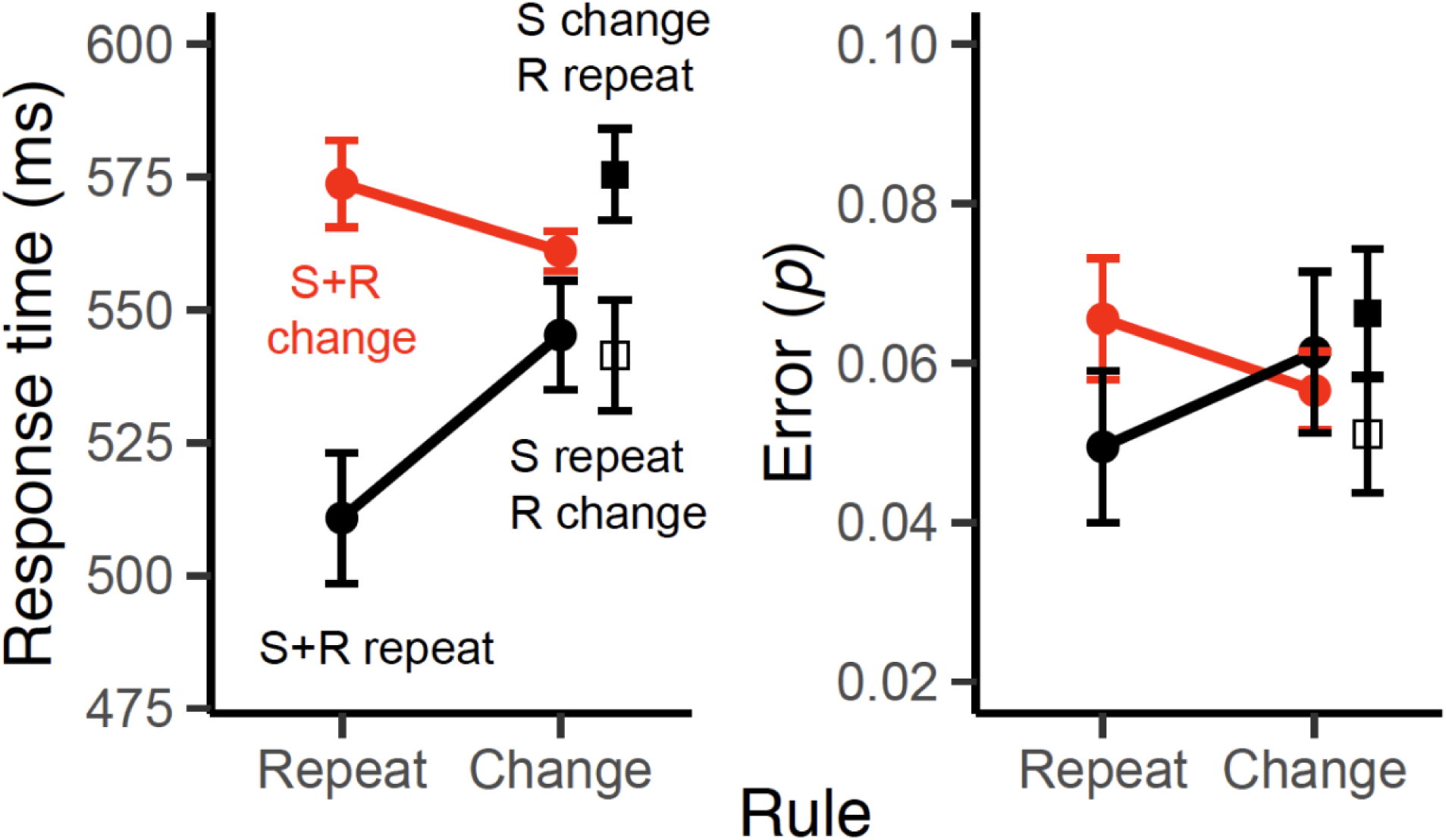
Mean RTs and errors for Experiments 2 as a function of the rule repetition/change, stimulus repetition/change, and the response repetition/change factors. Note that, in rule-repeat trials, partial updates of S-R settings (e.g., S change + R repeat) are not possible. Error bars specify 95% within-subject confidence intervals.

#### Decoupling Rule-specific and Rule-independent Conjunctions

We used the same analysis approach as in Experiment 1, only that here we included RSA models for both rule-specific (rule S-R) conjunctions and rule-independent (S-R) conjunctions. We found evidence that both types of conjunctions emerged over and above the constituent features (Fig. 7a). The rule-S-R conjunctions became active right after stimulus onset and were sustained robustly during the selection period. Again, activation of these representations preceded the emergence of response information (Supplementary Fig. 7). In contrast, the rule-independent, S-R conjunctions appeared immediately after rule-S-R conjunctions, but remained relatively weak compared to other action representations. We also replicated the pattern of results from Experiment 1 for the constituent features, with the exception that the rule representations diminished after stimulus onset (Fig. 7a). Excluding conjunction models restored the post-stimulus rule representation, suggesting that the rule S-R conjunction model captures the same variance as the rule model explains in this phase of action selection (Fig. 7a inset).

**Fig. 7.**
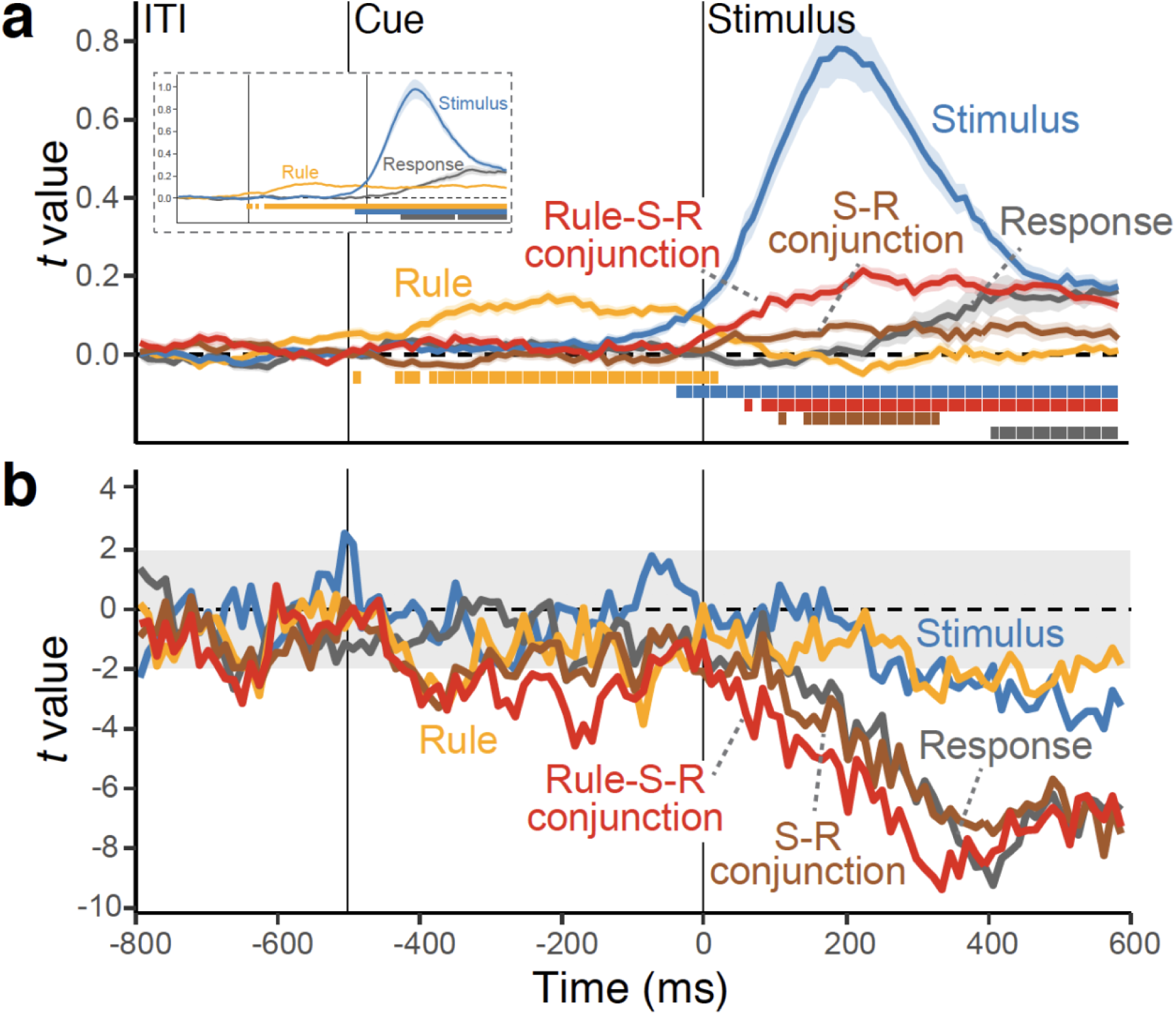
**a**, Average, single-trial *t*-values associated with each of the basic features and their conjunctions, derived from the RSA analysis (see Fig. 1c and 5). Shaded regions show the standard error around the mean. The colored squares at the bottom of the figure denote the significant time points using a non-parametric permutation test. The insert shows the same RSA fit scores when the conjunctions (i.e., rule S-R conjunction model and S-R conjunction model) were not included as predictors in the RSA analysis. **b**, Time-course of *t* values from multilevel, linear models predicting the variability in trial-to-trial RTs (the “impact” of representations on behavior), using RSA scores of all features as the simultaneous predictors. RSA model vectors for stimulus, response, rule, and conjunction representations. RSAs were performed separately within a subset of action constellations (i.e., G1 and G2) to orthogonalize all features (see the Method and Fig. 5ab for details).

Next, we examined again, which representations were the main driver of action selection. As shown in Fig. 7b, both rue-S-R conjunctions and S-R conjunctions explained substantial, and independent variability in trial-to-trial RTs, over and above the variance explained by the constituent features (Table 2 for the statistical results and Supplementary Fig. 4 for results from standard decoding analyses). These results replicate the findings from Experiment 1 that conjunctions are indeed critical of efficient action selection. In addition, they clarify that both rule-independent and rule-specific conjunctions are about equally important in predicting behavior, with possibly a slight edge for the rule-specific conjunctions.

**Table 2.** Predicting trial-by-trial RTs using the strength of decoded representations.

| Decoded Variable | Pre-stimulus |  | Early Post-stimulus |  | Late Post-stimulus |  |
| --- | --- | --- | --- | --- | --- | --- |
|  | *b* (*se*) | *t*-value | *b* (*se*) | *t*-value | *b* (*se*) | *t*-value |
| Rule | -.019 (.005) | -3.80 | -.017 (.013) | -1.34 | -.007 (.003) | -2.54 |
| Rule S-R Conjunction |  |  | -.061 (.012) | -5.00 | -.019 (.002) | -8.70 |
| S-R Conjunction |  |  | -.053 (.014) | -3.87 | -.022 (.003) | -7.32 |
| Stimulus |  |  | -.010 (.007) | -1.42 | -.008 (.002) | -3.94 |
| Response |  |  | -.037 (.014) | -2.57 | -.016 (.002) | -6.60 |

#### Conjunctive Representations and Partial-Overlap Costs

As in Experiment 1, we also examined the relationship between conjunction codes and behavioral partial-overlap costs. We found that only the strength of rule-S-R conjunctions showed the partial-overlap costs, *b*=-.021, *SE*=.009, *t*(21)=-2.22, for the early selection phase; *b*=-.021, *SE*=.009, *t*(21)=-2.24, for the late selection phase (Fig. 8a). None of the constituent features, *t*(21)>-.72), or S-R conjunctions showed such an effect, *b*=-.012, *SE*=.009, *t*(21)=1.27 for the early selection phase; *b*=-.007, *SE*=.010, *t*(21)=-.72, for the late selection phase. In addition, the strength of late rule-S-R conjunctions on the previous trial again significantly modulated RT partial-overlap costs on the next trail (Fig. 8b), *b*=.031, *SE*=.011, *t*(20)=2.81. This pattern was absent for constituent features, all *t*(21)<.38, and S-R conjunctions, *b*=-.009, *SE*=.011, *t*(21)=-.85. Thus, only the conjunctions that integrate rule information show a tight relation with the main behavioral indicator of event files, the partial-overlap cost.

**Fig. 8.**
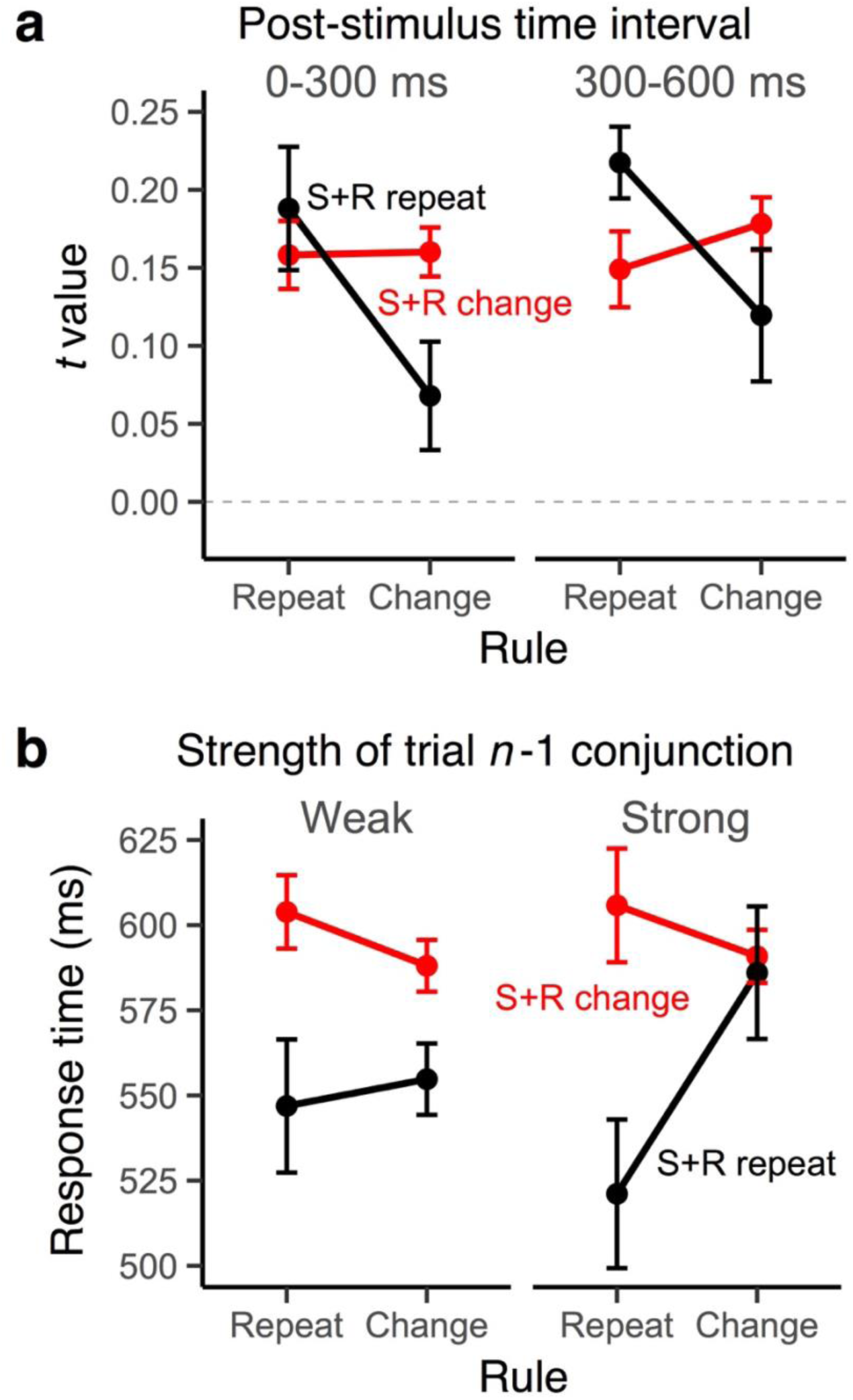
**a**, Average RSA scores of the rule-S-R conjunction as a function of the rule repetition/change and the stimulus-response repetition/change factors for early (0-300 ms) and late (300-600 ms) periods in the post-stimulus interval. **b**, Modulation of partial-overlap priming patterns in trial *n* as a function of the strength of the rule S-R conjunction in trial *n* −1 (median split). Error bars specify 95% within-subject confidence intervals.

## Discussion

We tested whether integrated, conjunctive representations between task-relevant features emerge during action selection, as postulated by event-file theory^7, 8^. In our paradigm action settings had to be updated flexibly for each trial, creating unique constellations between rules, stimuli and responses. We combined a standard, linear decoding approach with a subsequent, time-resolved RSA in order to track the emergence of conjunctive representations and their constituent features over time, and for each individual trial. In Experiment 1, conjunctions could entail any pairwise, or complete combination of rule, stimulus, or response features; in Experiment 2, we were further able to dissociate between rule-S-R conjunctions, and rule-independent S-R conjunctions.

The time course of decoded information showed a highly plausible cascade of action representations (rule, stimulus, and then response), and most critically, we found robust evidence for conjunctive representations—emerging shortly after stimulus onset and then persisting until response execution. Analyses with response-locked EEG data fully confirmed this pattern of results (Supplementary Fig. 7). The fact that conjunctive representations are continuously present from stimulus processing to response execution is consistent with their role in translating perceptual codes into response codes based on the current task rules. Even though the strength of conjunctive representations was on average much weaker than that of the constituent features, they were highly robust and consistent within individuals (Fig. 3 and Fig. 7 and Supplementary Fig. 9). Even more importantly, conjunctive representations were strong and unique predictors of trial-by-trial variability in RTs, over and above other constituent features. These results are difficult to reconcile with traditional stage theories^1–5^, where information flows in a strictly feed-forward manner and therefore does not allow the emergence of integrated representations. These results are also inconsistent with hierarchical control models that assume independent selection processes on different hierarchical levels. ^12, 13^ Instead, our results indicate that action selection is established by tying together disparate, task-relevant features from the entire selection event into a common representation.

The fact that in Experiment 2 rule-S-R conjunctions *and* rule-independent, S-R conjunctions emerged is an important result in its own right. It suggests that integrated representations that match the contingencies in the environment can develop in parallel, and on different levels of abstraction. This combination of both highly specific and rule-general representations can account for the fact that S-R associations learned within one rule can transfer to another rule, albeit in a limited manner.^10, 21^ It is also consistent with the proposal that event files themselves can possess an internal, hierarchical organization.^9^

A key behavioral indicator of event files is the partial-overlap priming pattern, which entails benefits when all action features either repeat or change and costs when there is partial overlap of features across trials.^10, 11^ In both experiments, we found that this pattern not only in RTs and errors (Fig. 2 and Fig. 6), but also in the strength of conjunctions (Fig. 4a and Fig. 8a). Even more importantly, the strength of conjunctions on trial *n*, predicts the size of partial overlap costs on trial *n*+1 (Fig. 4b and Fig. 8b), suggesting the stronger action features are tied together into conjunctions, the harder it is to “unbind” them on the following trial in order to integrate them into a new conjunction. Recent behavioral studies have raised questions about whether the strength of partial-overlap costs is explained by the strength of the initial binding, or instead by difficulty of selectively retrieving integrated, action-relevant features.^8^ While the present results do not rule out the contribution of retrieval-related effects, they do point to the “binding strength” of the original conjunction as a critical factor that determines partial-overlap costs.

It is particularly important that the tight relationship with the partial-overlap pattern was only found for conjunctions (i.e., rule-S-R conjunction in Experiment 2), thus functionally dissociating conjunctions from their constituent codes. Moreover, the results in Experiment 2 also indicated that only rule-specific S-R conjunctions were related to the partial-overlap cost, not however the rule-independent S-R conjunctions. As noted, for Experiment 1, the task design did not allow firm conclusions about whether or not conjunctions contained rule-specific information. However, the conjunctions in Experiment 1 showed a similar priming pattern as the rule-S-R conjunctions in Experiment 2, suggesting integration of not just stimuli and responses, but also of rules in both experiments.

While our results indicate with high resolution when representations of specific features and feature combinations are activated, they provide no neuroanatomical information (see Supplementary Results). Cell-physiological work with monkeys and human, neuroimaging work indicates that the representation of task-relevant features, including rules, is distributed across large areas frontal and parietal cortex.^22, 23^ From animal models, there is substantial evidence that the hippocampus and the frontal cortex are particularly important for representing conjunctive information.^24, 25^ Human neuroimaging work also mainly implicates the hippocampus ^26, 27^; attempts to decode task representations in the frontal areas have proven more challenging^28^, but have also seen some recent success.^29, 30^

In nonhuman primates, single cell recordings have also shown that while basic task features (cues, rules, stimuli, and responses) are encoded across various frontal and parietal areas during rule-based action selection^22, 31^, a substantial proportion of recorded neurons are tuned to the mixture of multiple features in a non-linear manner.^24, 32^ Such heterogeneous, neural responses allow both efficient, linear read-out of information to downstream neurons and can also code high-dimensional, conjunctive information^33^.

An important finding from this research is that the degree of conjunctive information coded in recorded neurons is functionally distinct from the representation of linear features. For example, high-dimensional, non-linear information was found to be highly robust on correct trial, but was largely missing on error trials, whereas low-dimensional information is equally strong on correct and error trials.^24^ This pattern is consistent with our finding that the strength of conjunctive representations uniquely predicts trial-by-trial performance, beyond the predictive strength of constituent, simple features (Fig. 3b and Fig. 7b). Further evidence for a functional dissociation comes the finding that conjunctive representations express trial-to-trial transitions (i.e., the partial-overlap priming pattern) in a qualitatively different manner than the constituent feature representations (see Fig. 4 and Fig. 8).

These results about the relevance of conjunctions for efficient action selection and the mismatch priming pattern also directly confirm predictions from event-file theory. Therefore, they provide an important, missing link between two, so-far distinct lines of research: The relatively abstract, event-file theory, designed to explain the architecture of human action selection, and the recent advances from animal research about the neural implementation of high-dimensional, non-linear representations. Beyond the current demonstration of the role of conjunctive representation in human action control, there is a range of important, open questions. For example, we do not know how these representations are constrained by capacity limitations^34^, to what degree they allow integration of action outcomes or goals^35^, or how they change through experience (see Supplemental Fig. 9).^10^ The decoding approach used here, promises answers to these and related questions.

## Method

### Participants

A total of 44 people participated after signing informed consent following the protocol approved by the University of Oregon’s Human Subjects Committee in exchange for the compensation of $10 per hour and the additional performance-based incentive. Participants with excessive amount of EEG artifacts (more than 35% of trials) were removed from further analysis. As a result, we retained 20 out of 22 participants for Experiment 1 and 21 out of 22 for Experiment 2.

### Stimuli, Tasks and Procedure

Participants performed a cued rule-selection task, in which one of the pre-instructed action rules, on trial-by-trial basis, was randomly selected to determine possible S-R mappings^10^; Fig. 1b). Based on the cued rule, participants responded to the location of a circle (1.32° in radius) that randomly appeared in the corner of a white frame (6.6° in one side) by selecting one of the four response keys that were arranged in 2 x 2 matrix. Each action rule specified four S-R mappings using a simple spatial transformation rule. For instance, the vertical rule mapped the left-top circle to the bottom-left response as a correct response and vice versa. We used two cues for each rule (a pair of verbal cues in Experiment 1 and symbol/word pair in Experiment 2) that appeared in either even or odd trials to prevent immediate cue repetitions. Thus, cues, rules, responses, and stimuli were orthogonalized, and the combination of these features generated unique action constellations.

In Experiment 1,“vertical”, “horizontal” and “diagonal” rules were randomly cued (i.e., 66.6 % switch rate). In Experiment 2, “vertical”, “horizontal”, “clockwise” and “counterclockwise” rules were used (i.e., 75% switch rate; Fig. 2c). Here, half of S-R links were shared across rules (e.g., a left-top circle leads to a left-bottom response in both the vertical and the counterclockwise rule). This allowed us to generate transitions between trials with rule changes but repetitions of S-R links (Fig. 1c).

There were two practice blocks and 200 experimental blocks in both studies. Participants were instructed to respond as fast and accurately as possible to complete as many trials as possible within each 16-second block. Trials that began within the 16 seconds were allowed to complete. Participants were given a performance-based incentive for trials with RTs faster than the 75th percentile of correct responses in the preceding blocks when 1) the overall accuracy was above 90 percent and 2) there were more than 7 completed trials in a given block. While performing the task, participants were asked to rest the index finger of their dominant hand in the center of the four keys in matrix and to hit the correct key. All stimuli were created in Matlab (Mathworks) using the Psychophysics Toolbox ^36, 37^ and were presented on a 17-inch CRT monitor (refresh rate: 60 Hz) at a viewing distance of 100 cm.

### EEG recordings and preprocessing

Electroencephalographic (EEG) activities were recorded from 20 tin electrodes held in place by an elastic cap (Electrocap International) using the International 10/20 system. The 10/20 sites F3, Fz, F4, T3, C3, CZ, C4, T4, P3, PZ, P4, T5, T6, O1, and O2 were used along with five nonstandard sites: OL halfway between T5 and O1; OR halfway between T6 and O2; PO3 halfway between P3 and OL; PO4 halfway between P4 and OR; and POz halfway between PO3 and PO4. Electrodes placed ~1cm to the left and right of the external canthi of each eye recorded horizontal electrooculogram (EOG) to measure horizontal saccades. To detect blinks, vertical EOG was recorded from an electrode placed beneath the left eye and reference to the left mastoid. The left-mastoid was used as reference for all recording sites, and data were re-referenced off-line to the average of all scalp electrodes. The EEG and EOG were amplified with an SA Instrumentation amplifier with a bandpass of 0.01–80 Hz and were digitized at 250 Hz in LabView 6.1 running on a PC.

EEG data was first epoched by 18 second intervals to include all trials within a block. After time-frequency decomposition was performed (see *Time-Frequency Analysis* section), these epochs were further segmented into trial-to-trial epochs (−600 ms to 600 ms intervals for Experiment 1 and −800 ms and 600ms intervals for Experiment 2, relative to the onset of a stimulus). These trial-to-trial epochs including blinks (>80uv, window size = 200 ms, window step = 50 ms), large eye movements (>1°, window size = 200 ms, window step = 10ms), blocking of signals (range = −0.01 uv to 0.01 uv, window size = 200 ms) were excluded from subsequent analyses. For all EEG analyses, error trials, post-error trials and trials with exceedingly slow RTs (i.e., slower than 99.5 % of all responses) were excluded to be consistent with behavioral analyses.

#### Time-Frequency Analysis

Temporal-spectral profiles of single-trial EEG data were obtained via complex wavelet analysis^38^ by applying time-frequency analysis to preprocessed EEG data segmented for each block (>18 seconds to exclude the edge artifacts). The power spectrum was convolved with a series of complex Morlet wavelets (*e*^2*πft*^*e*^−*t*2/(2*σ2)^), where *t* is time, *f* is frequency increased from 1 to 35 Hz in 35 logarithmically spaced steps, and σ defines the width of each frequency band, set according to *n/2f*, where *n* increased from 3 to 10. The logarithmic scaling was used to keep the width across frequency band approximately equal, and the incremental number of wavelet cycles was used to balance temporal and frequency precision as a function of frequency of the wavelet. After convolution was performed in the frequency-domain, we took an inverse of the Fourier transform, resulting in complex signals in the time-domain. A frequency band-specific estimate at each sample point was defined as the squared magnitude of the convolved signal *Z*(real([*z*(*t*)]2 + imag[*z*(*t*)]2) for instantaneous power.

### Representational Similarity Analysis

Our goal was to obtain information about the strength of each feature and conjunction on the level of individual trials and timepoints within trials. This required a two-step procedure. First, we performed a linear decoding analysis to discriminate between all 12 different action constellations in Experiment 1, or 16 constellations in Experiment 2. Specifically, we performed a penalized linear discriminant analysis using the caret package in R^39–41^. At every time sample point, the power of rhythmic EEG activity was averaged within the predefined ranges of frequency values (1-3 *Hz* for the delta-band, 4-7 *Hz* for the theta-band, 8-12 *Hz* for the alpha-band, 13-30 *Hz* for the beta-band, 31-35 *Hz* for the gamma-band), generating 100 features (5 frequency-bands X 20 electrodes) to train decoders. Within individuals, these data points were z-transformed across electrodes at every sample to remove the effects that uniformly influenced all electrodes. We used a *k*-fold repeated cross-validation procedure to evaluate the decoding results^42^, by randomly partitioning single-trial EEG data into four independent folds. The number of observations of each action constellation was kept equal within and across folds by dropping trials randomly. Three folds served as a training set and the remaining fold was used as a test set; this step was repeated until each fold served as a test set. Each cross-validation cycle was repeated eight times, in which each step generated a new set of randomized folds. Resulting classification probabilities (i.e., evidence estimated for each case of S-R mapping) were averaged across all cross-validated results with the best tuned penalty parameters. This decoding step yielded a vector of “confusion profiles” of classification probabilities for both the correct and all possible incorrect classifications and for each time point and trial (Fig. 1c).

As a second step, we then applied RSAs^17^ to each profile of classification probabilities in order to determine their underlying similarity structure for each time point and trial. Specifically, we regressed the confusion vector onto model vectors as predictors, which were derived from a set of representational similarity model matrixes. Each model matrix uniquely represents a potential, underlying representation (e.g., rules, stimuli, responses and conjunctions; Fig. 1c and Fig. 5b). For example, the rule model predicts neural responses to be similar (i.e., more confusable) among instances of the same rule, but dissimilar across different rules. To estimate the unique variance explained by competing models, we regressed all model vectors simultaneously. Thus, we obtained coefficients for each of the four model vectors (e.g., rule, stimulus, response, conjunction for Experiment 1). These coefficients (i.e., their corresponding *t*-values) allowed us to relate the dynamics of action representations to trial-to-trial variability in behavior (see *Multilevel Modeling* section for details). In all RSAs, we logit-transformed classification probabilities and further included a subject-specific “conjunction RT” model (i.e., a vector of z-scored, RTs, averaged for each subject and action constellation) as a nuisance predictor to reduce potential biases in decoding due to idiosyncratic differences in RTs among action constellations. We excluded resulting *t*-values that exceeded 5 SDs from means for each sample point, which excluded 0.12% and 1.32% of the entire samples Experiments 1 and 2 respectively. Resulting *t-*values were averaged within in 12 ms non-overlapping time samples.

In Experiment 1, we constructed RSA models for the rules, stimuli, responses, and conjunctions (Fig. 1c). In Experiment 2, the conjunction model was separated for the rule-specific S-R conjunction model (rule-S-R conjunction) and the rule-independent S-R conjunction model (S-R conjunction; Fig. 5b). Complete orthogonalization of features could be established within each of two equal-sized subspaces of the entire space of action constellations, but not across the entire space. Therefore, we performed the RSA within each of these subspaces independently and subsequently averaged the results. Specifically, one subspace (G1 in Fig. 5) contained constellations with stimuli at the top-left or bottom-right corner (leading to a bottom-left or bottom-right response for all rules), whereas the second subspace (G2 in Fig. 5) contained trials with stimuli at the left-bottom or top-right corner (leading to a top-left or bottom-right response). Within each subspace, conjunctions were defined by the combination of four rules (vertical, horizontal, clockwise, and counterclockwise), two stimulus positions, and two responses, ensuring that each S-R link could occur in the context of two different action rules.

#### Non-Parametric Permutation Test

To test statistical significance of all time-resolved decoding results (Fig. 3a and Fig. 7a and Supplementary Fig. 3 and Supplementary Fig. 4) while accounting for multiple comparisons, we carried out nonparametric permutation tests using the single-threshold method^43^. For each feature, we computed permutation distributions of the maximum statistic for every sample point from −200 ms prior to the onset of the cue to 600 ms after the onset of the stimulus. Specifically, we first obtained classification results (and performed RSA for Fig. 3a and Fig. 7a) by decoding of data with randomly shuffled condition labels. We then performed a series of *t*-tests for every sample against the null level (i.e., the chance level). For the RSA results, the null level was 0 for *t*-values. Out of the series of *t*-test results, we retained the maximum *t*-value. We repeated this process 10000 times by randomly drawing samples from all possible permutations of labels, thereby generating the permutation distributions of the maximum statistics. This approach allowed us to identify statistically significant time points by comparing scores from the correct labels to the critical threshold, which was defined as the 99th (i.e., alpha =.01) of the largest member of maximum statistics in the permutation distribution of the corresponding variable.

### Multilevel Modeling

We used multilevel linear modeling to analyze trial-by-trial variability in decoded representations and their relationship to RTs. The models estimated fixed effects of predictors as well as subject specific intercepts and slopes as random effects. For all statistical tests, the dependent variable (e.g., RSA scores or RTs) was prewhitened by the linear and quadratic trends of experimental trials and blocks. RTs were further log-transformed before the fitting. We performed statistical tests for a-priori selected time intervals: cue-to-stimulus period from the onset of cue to the onset of stimulus (−300 to 0 ms for Experiment 1 and −500 to 0 ms for Experiment 2), early post-stimulus period (0 to 300 ms of the post-stimulus segment for both studies), and late post-stimulus period (300 to 600 ms of the post-stimulus segment for both studies). We predicted trial-to-trial RTs/RSA scores in the current trials with EEG signals from pre-stimulus and early post-stimulus periods in hopes of capturing processing prior to response execution (see also Supplementary Fig. 7 for results using signals aligned to the response onsets). The late post-stimulus interval was used to assess how partial-overlap costs are modulated by the strength of action representations developed during selection in *n*−1 trials. In addition, we separately performed a series of regressions to visualize changes in RT predictability—”impact” of moment-to-moment strength of decoded feature—by fitting models at each sample point without random slopes (Fig. 3b and Fig. 7b).

## Acknowledgements

Acknowledgements: This work was supported in part by National Institute of Aging grant R01 AG037564-01A1, by NSF grant 1734264 and an Award by the Humboldt Foundation to Ulrich Mayr

## Supplementary Results

### Decoding of basic, constituent features without RSA

In the main paper, we used RSA analyses to distinguish conjunction representations from the representation of constituent features. In order to compare these results with a standard decoding approach, we also performed standard multivariate decoding analyses for each constituent feature independently^1^. The analysis procedure (i.e., cross-validation, non-parametric permutation test, and subsequent multilevel modeling predicting trial-to-trial RTs) was identical to the method for RSA except for the following points: 1) final outputs of decoding analysis were classification probabilities (then logit-transformed) rather than RSA fit score, and 2) individuals-specific mean RTs of all action constellations were included as a control predictor in multilevel models of RTs.

Supplementary Fig. 3 and Supplementary Fig. 4 show the trajectories of classification probabilities of constituent features (rules, stimuli and responses) and their impact on trial-to-trial variability in RTs (see the inserts of Fig. 3a and Fig. 6a for the corresponding RSA results). The results were overall consistent with RSA results, when excluding the conjunction models (i.e., see inserts for Fig. 2a and Fig. 4a). The rule was activated during the pre-stimulus phase, followed by a strong expression of the stimulus, and finally by the response. This pattern directly replicates results using a more standard task-switching paradigm^1^. These results also confirmed that our RSA approach produced qualitatively similar results to the standard time-resolved decoding analysis^2^ for constituent features (when the conjunction model was excluded).

### RSA using frequency-specific EEG activity

Previous studies showed that specific control representations are encoded in the frequency-specific, rhythmic EEG activity (e.g., the ordinal position codes in the theta-band (4-7 *Hz*)^3^. As an exploratory analysis, we analyzed how different frequency-bands contribute to the decoding of both conjunctions and constituent features. For this, we replicated the combination of decoding and RSA analyses separately for the delta (1-3 *Hz*), theta (4-7 *Hz*), alpha (8-12 *Hz*), beta(13-30 *Hz*) and gamma (31-35 *Hz*) frequency bands. To reduce the influence of temporal smearing, which could differ across frequency-bands^4^, we averaged data over a-priori selected time intervals (i.e., pre-stimulus, early and late post-stimulus phase) prior to the training of decoders. Other steps in the analysis were identical to the one described in *Representational Similarity Analysis* section.

Supplementary Fig. 5 and Supplementary Fig. 6 summarize RSA scores of different action features among individuals for each time interval, compared to the results using all five frequency-bands. Overall, participants exhibited considerable variability in terms of the frequency-bands in which specific information was expressed. However, the following points seem to be consistent across individuals: 1) stimulus positions in early post-stimulus interval were coded in the theta and the alpha-band, a finding that is consistent with earlier decoding results^5^, and 2) conjunctions (i.e., rule-S-R conjunctions in Experiment 2) tended to be expressed most strongly in the delta band. The fact that the neighboring theta-band activity only weakly contributed to the conjunctions emphasizes the special role of delta-band activity. We had no a-priori predictions about frequency-specific effects. However, we note that there are recent reports suggesting that delta-band frequencies carry information about abstract decision variables^6^. Using a similar “search-light” method across electrodes (instead of frequency-space), we also checked for relative contribution of local scalp regions for expression of representations that are more localize consistent across individuals. However, we found the patterns with which representations were expressed to be highly idiosyncratic (see Kikumoto and Mayr^3^ for similar results in hierarchical serial-control task).

### RSA time-aligned to responses

The trajectory of RSA fit scores, summarized in Fig. 3a and Fig. 6a, revealed how action-relevant representations unfold in reference to the onset of stimulus. However, these decoding results contain a limited number of samples which responses were executed during the post-stimulus interval (average RT: *M* = 490 ms, *SD* = 1.46 ms in Experiment 1; *M* = 490 ms, *SD* = 1.67 ms in Experiment 2). To ensure that our stimulus-locked decoding results are not affected by such early responses, and to provide information about how representations unfold relative to responses, we performed an additional RSA using EEG data that are aligned to the onset of each trial’s response. Supplementary Fig. 7 shows the corresponding time-course of RSA scores for Experiments 1 and 2. These results are highly consistent with stimulus-locked results in showing early peaks for rule, followed by stimulus, then conjunction, and finally response representations, which peaked just before responses were executed.

#### Cross-session RSA

We know little about within-subject consistency of EEG-decoded, complex action representations across longer temporal intervals. In particular, given that the neural patterns that we decoded were highly idiosyncratic (see Section *RSA using frequency-specific EEG activity*), it is important to explicitly test the stability of such patterns within individuals. We were able to recruit 9 out of the 20 participants from Experiment 1 for a second session, 6 months after the first session.

Participants responded significantly faster in the second session than in the first session, RTs: *F*(1,8)=29.05, *MSE*=735.54, *p*<.001,ηp2=.78; errors: *F*(1,8)=.40, *MSE*<.001, *p*=.54, ηp2=.05 (Supplementary Fig. 7). In order to perform cross-session RSAs we trained decoders using data within each session separately. Supplementary Fig. 9 (left column) summarizes differences in RSA scores of each representation across sessions. Both the pre-stimulus rule representation and the conjunction in the stimulus-to-response phase were enhanced in the later session. Then, decoders that were trained separately within sessions were applied to EEG data of the other session at matching time points. As shown in Supplementary Fig. 9 (right column), all features, including the conjunctive representation, showed robust generalizability across sessions. This pattern indicates that the neural representation of action-relevant representations is remarkably stable. An important implication of this result is that in future research such techniques can be applied to analyze how action-relevant representations are shaped through experience or consolidation.

## Supplementary Figures

**Fig S1.**
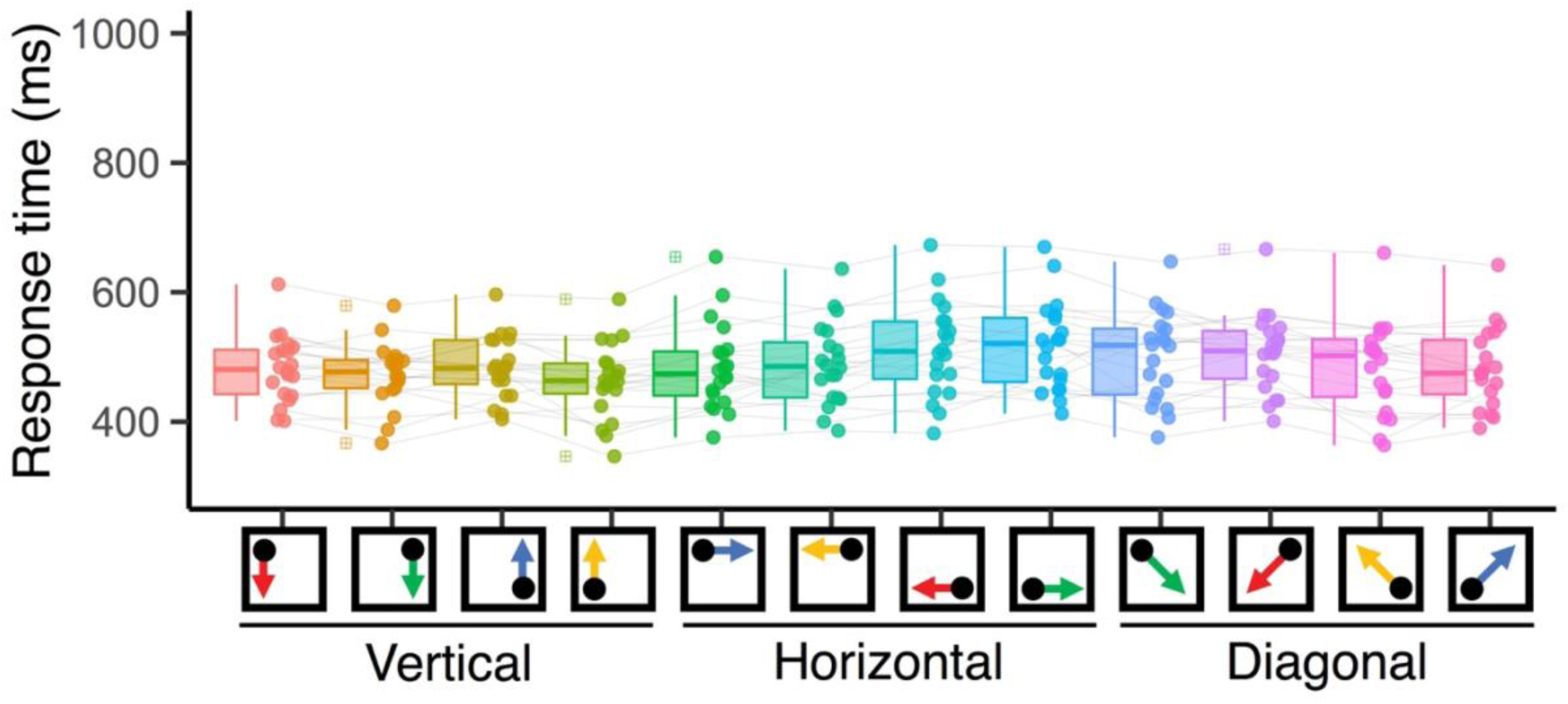
Mean RTs of individual subjects for all action constellations in Experiment 1. Subjects-specific RT vectors were included as a nuisance predictor during RSA fitting.

**Fig. S2.**
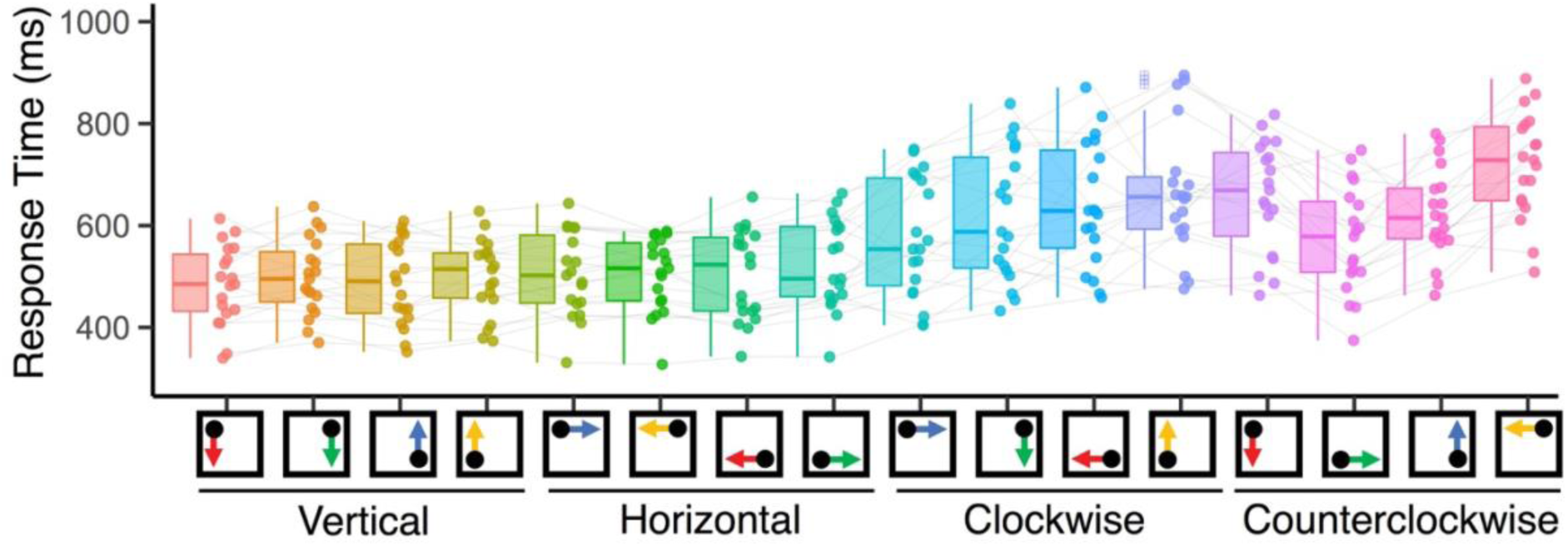
Mean RTs of individual subjects for all action constellations in Experiment 2. Participants responded slowly for trials with the non-symmetric translation (clockwise and counterclockwise rules) compared to the symmetric rules (vertical and horizontal rules) as reported previously7. Subjects-specific RT vectors were included as a nuisance predictor during RSA fitting.

**Fig. S3.**
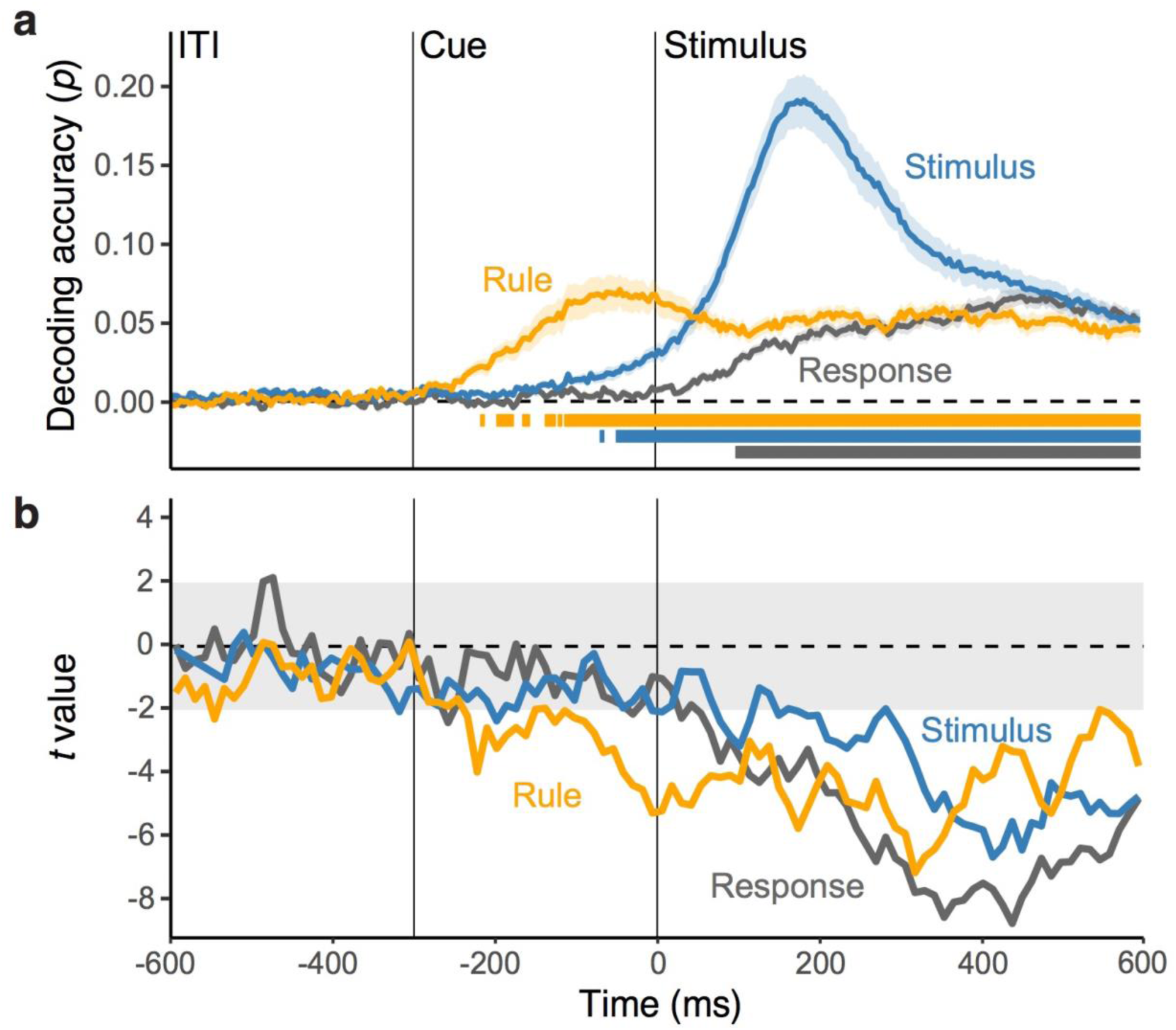
**a**, Average decoding accuracy of constituent features over time, derived from a standard decoding analysis in Experiment 1. Shaded regions specify the standard error around the mean. Squares below lines denote the significant time points correcting for multiple comparison using a non-parametric permutation test. **b**, Time-course of *t* values from multilevel, linear models predicting the variability in trial-to-trial RTs, using single-trial classification probability of each feature as predictors simultaneously.

**Fig. S4.**
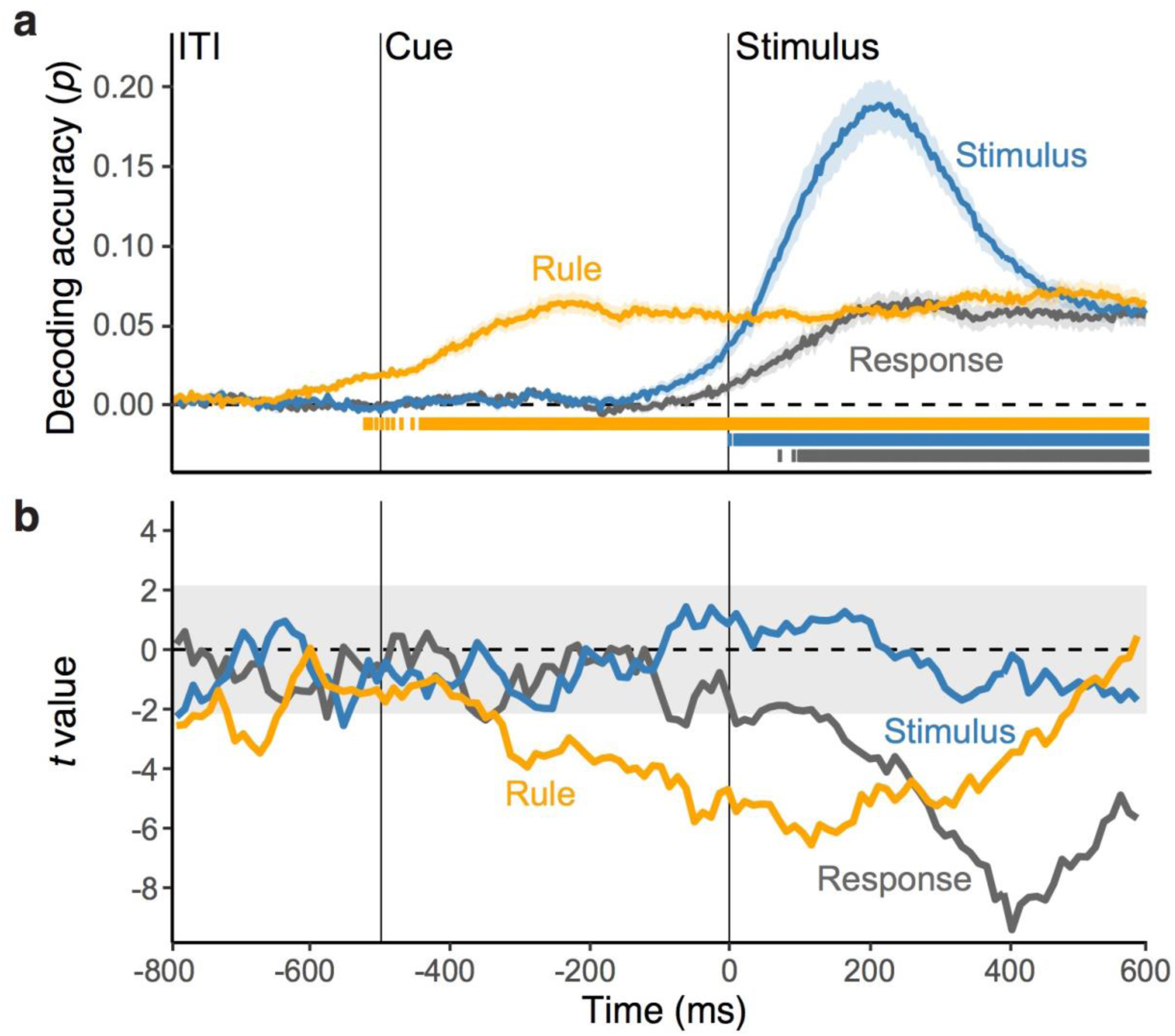
**a**, Average decoding accuracy of constituent features over time, derived from a standard decoding analysis in Experiment 2. Shaded regions specify the standard error around the mean. Squares below lines denote the significant time points correcting for multiple comparison using a non-parametric permutation test. **b**, Time-course of *t* values from multilevel, linear models predicting the variability in trial-to-trial RTs, using single-trial classification probability of each feature as predictors simultaneously.

**Fig. S5.**
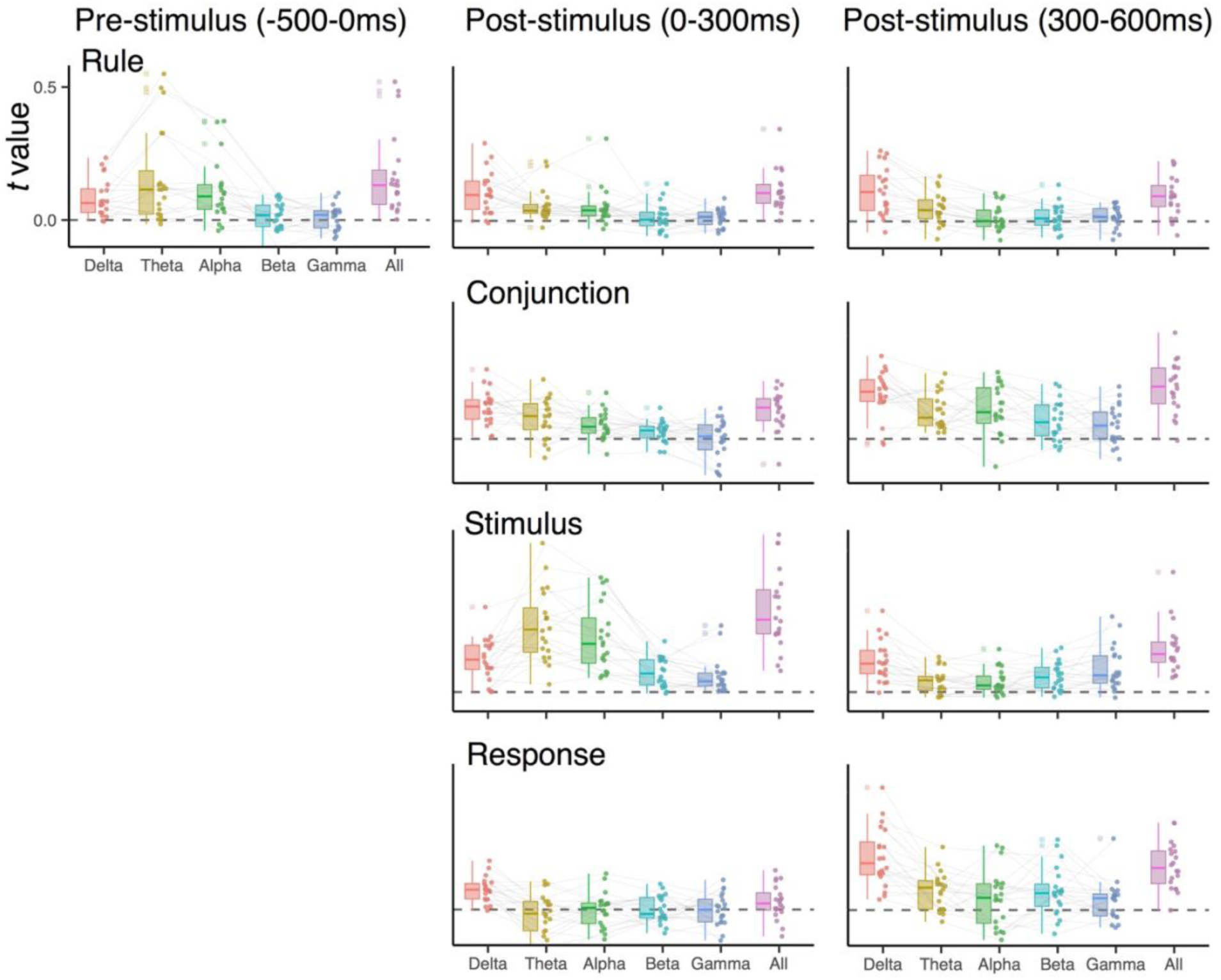
Experiment 1: RSA scores when decoding EEG signals in specific frequency ranges (1-3 *Hz* for the delta-band, 4-7 *Hz* for the theta-band, 8-12 *Hz* for the alpha-band, 13-30 *Hz* for the beta-band, 31-35 *Hz* for the gamma-band, and 1-35 *Hz* for all). EEG signals were averaged over pre-stimulus (−300 to 0 ms), early post-stimulus (0 to 300 ms) and late post-stimulus (300 to 600 ms) time intervals before the decoding analysis.

**Fig. S6.**
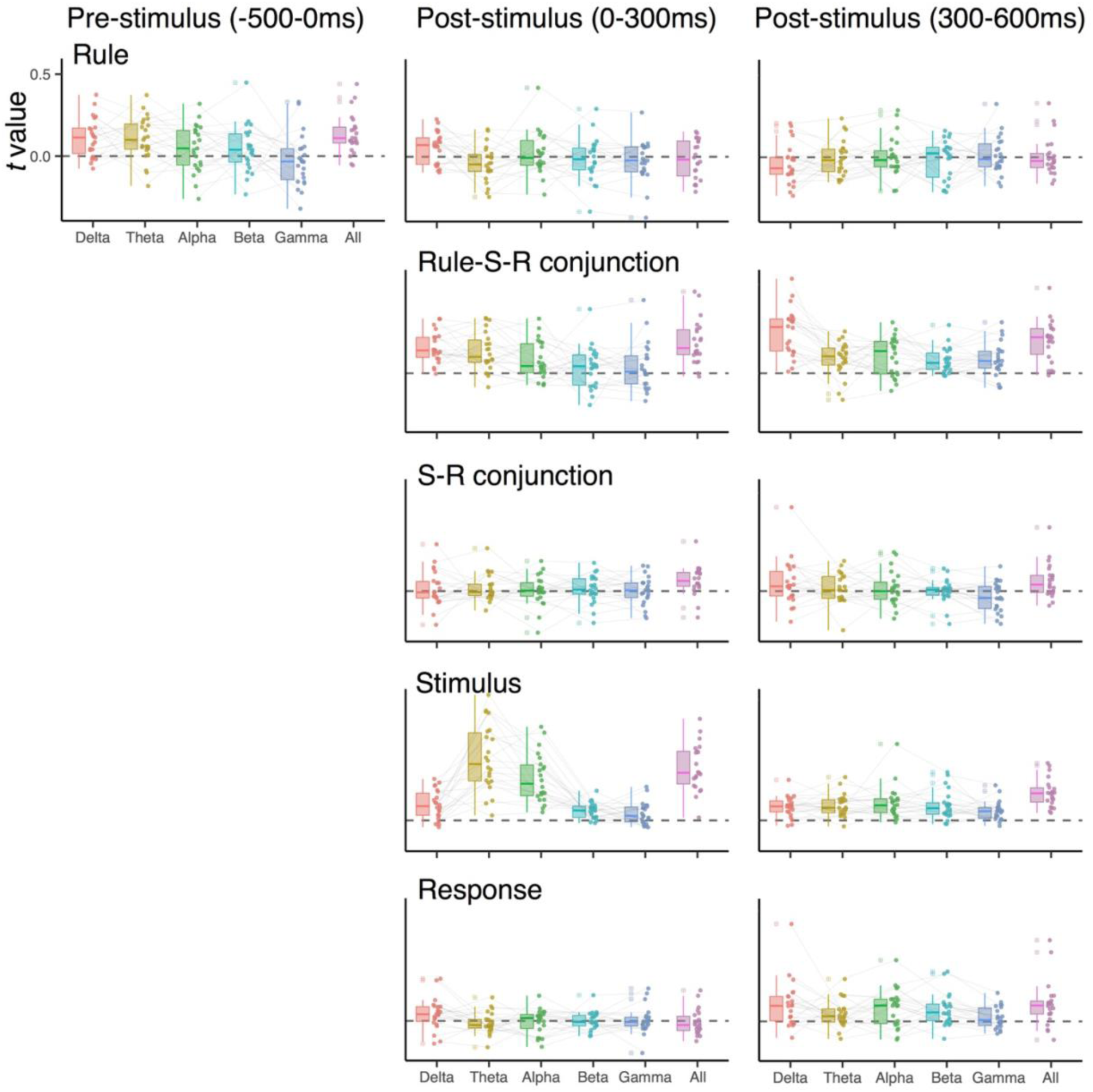
Experiment 2: RSA scores when decoding EEG signals in specific frequency ranges (1-3 *Hz* for the delta-band, 4-7 *Hz* for the theta-band, 8-12 *Hz* for the alpha-band, 13-30 *Hz* for the beta-band, 31-35 *Hz* for the gamma-band, and 1-35 *Hz* for all). EEG signals were averaged over pre-stimulus (−300 to 0 ms), early post-stimulus (0 to 300 ms) and late post-stimulus (300 to 600 ms) time intervals before the decoding analysis.

**Fig. S7.**
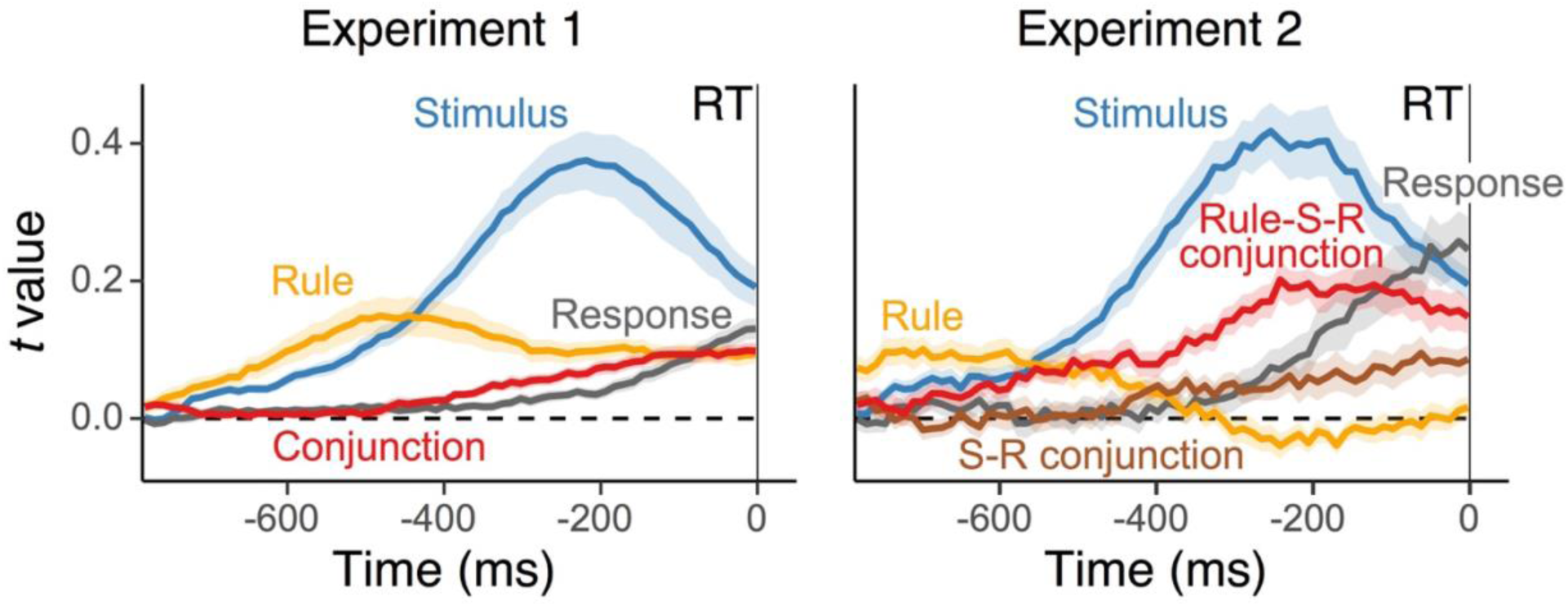
Average, single-trial RSA scores for each representation, using EEG signals aligned to the onset of trial-to-trial response events. Shaded regions specify standard error of the mean.

**Fig. S8.**
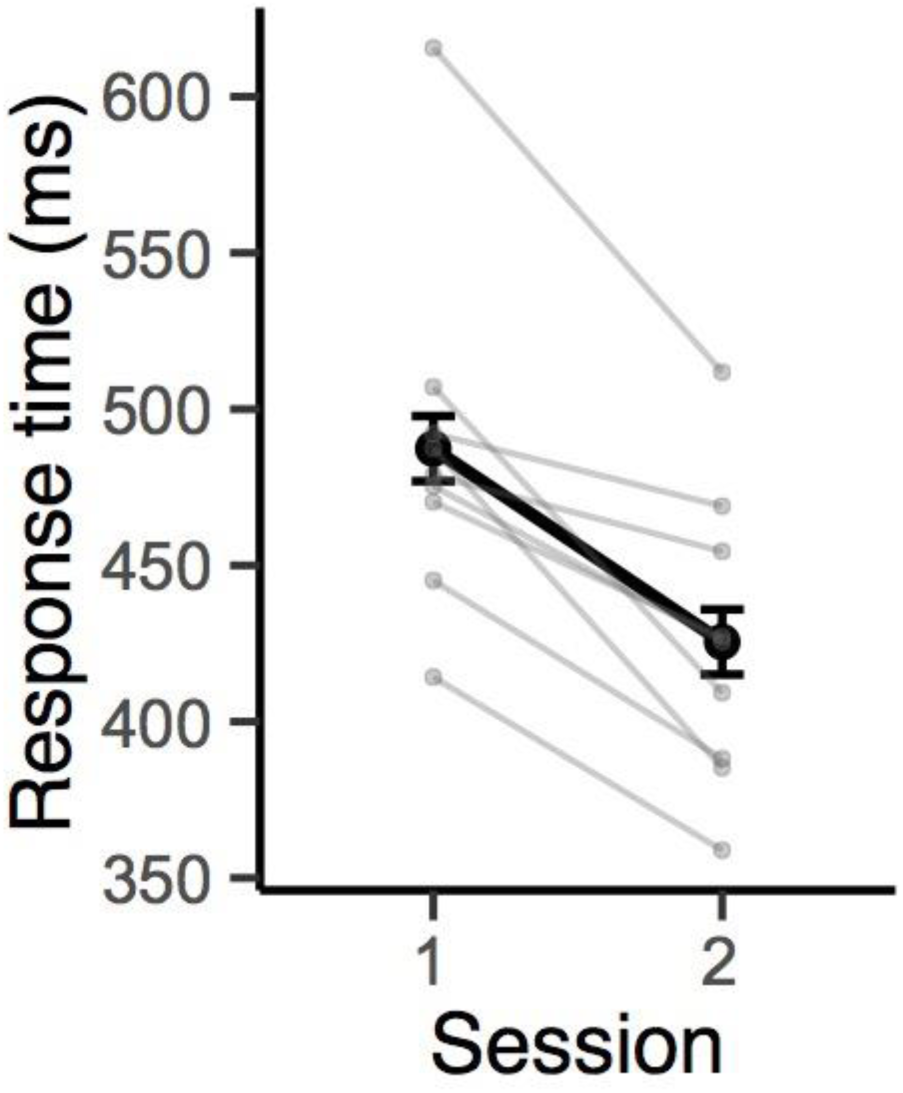
Individuals’ mean RTs and group average RTs (black) between the first session and the second session at least 6 months later (*n* = 9) for Experiment 1. Error bars specify 95% within-subject confidence intervals.

**Fig. S9.**
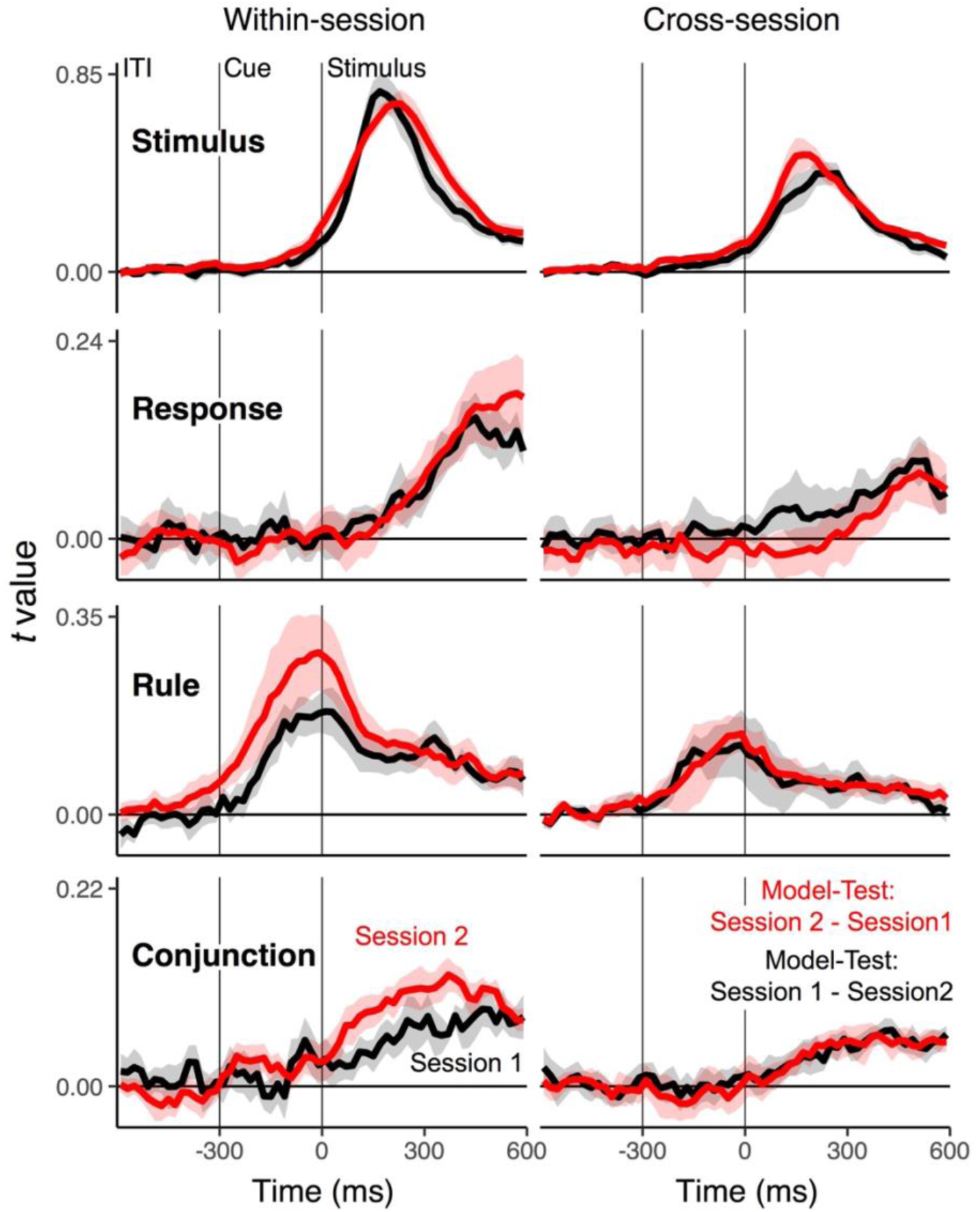
Average, single-trial RSA fit scores for conjunctions and constituent features across sessions (*n* = 9). *Left panels*: Time-course of RSA scores training decoders. *Right panels*: RSA scores from cross-decoding across sessions. Decoders are trained with EEG data from the session 1 (shown in red) or session 2 (shown in black) and applied on data from the counterpart session to test their generalizability. Shaded regions specify 95% within-subject confidence intervals.

## Supplemental Tables

**Supplementary Table 1.**
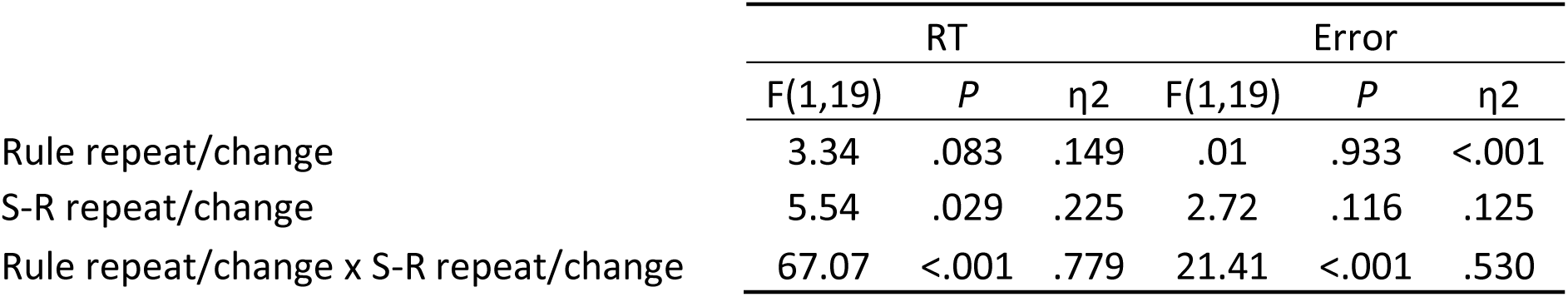
Anovas of RTs and errors with the factors rule repeat/change and S-R repeat/change

|  | RT |  |  | Error |  |  |
| --- | --- | --- | --- | --- | --- | --- |
| | F(1,19) | *P* | $\eta^2$ | F(1,19) | *P* | $\eta^2$ |
| Rule repeat/change | 3.34 | .083 | .149 | .01 | .933 | <.001 |
| S-R repeat/change | 5.54 | .029 | .225 | 2.72 | .116 | .125 |
| Rule repeat/change x S-R repeat/change | 67.07 | <.001 | .779 | 21.41 | <.001 | .530 |

**Supplementary Table 2.** Anovas of RTs and errors with the factors rule repeat/change and S-R repeat/change

|  | RT |  |  | Error |  |  |
| --- | --- | --- | --- | --- | --- | --- |
| | F(1,20) | P | $\eta^2$ | F(1,19) | P | $\eta^2$ |
| Rule repeat/change | 5.08 | .036 | .202 | .07 | .791 | .003 |
| S-R repeat/change | 42.28 | <.001 | .679 | 1.72 | .204 | .079 |
| Rule repeat/change x S-R repeat/change | 25.56 | <.001 | .561 | 6.91 | .002 | .257 |

